# BEGAN: Boltzmann-Reweighted Data Augmentation for Enhanced GAN-Based Molecule Design in Insect Pheromone Receptors

**DOI:** 10.1101/2024.09.07.611826

**Authors:** Jialei Dai, Yutong Zhang, Chen Shi, Yang Liu, Peng Xiu, Yong Wang

## Abstract

Identifying molecules that bind strongly to target proteins in rational drug design is crucial. Machine learning techniques, such as generative adversarial networks (GAN), are now essential tools for generating such molecules. In this study, we present an enhanced method for molecule generation using objective-reinforced GANs. Specifically, we introduce BEGAN (Boltzmann-Enhanced GAN), a novel approach that adjusts molecule occurrence frequencies during training based on the Boltzmann distribution exp(*−*Δ*U/τ*), where Δ*U* represents the estimated binding free energy derived from docking algorithms and *τ* is a temperature-related scaling hyperparameter. This Boltzmann reweighting process shifts the generation process towards molecules with higher binding affinities, allowing the GAN to explore molecular spaces with superior binding properties. The reweighting process can also be refined through multiple iterations without altering the overall distribution shape. To validate our approach, we apply it to the design of sex pheromone analogs targeting *Spodoptera frugiperda* pheromone receptor SfruOR16, illustrating that the Boltzmann reweighting significantly increases the likelihood of generating promising sex pheromone analogs with improved binding affinities to SfruOR16, further supported by atomistic molecular dynamics simulations. Furthermore, we conduct a comprehensive investigation into parameter dependencies and propose a reasonable range for the hyperparameter *τ*. Our method offers a promising approach for optimizing molecular generation for enhanced protein binding, potentially increasing the efficiency of drug discovery pipelines.

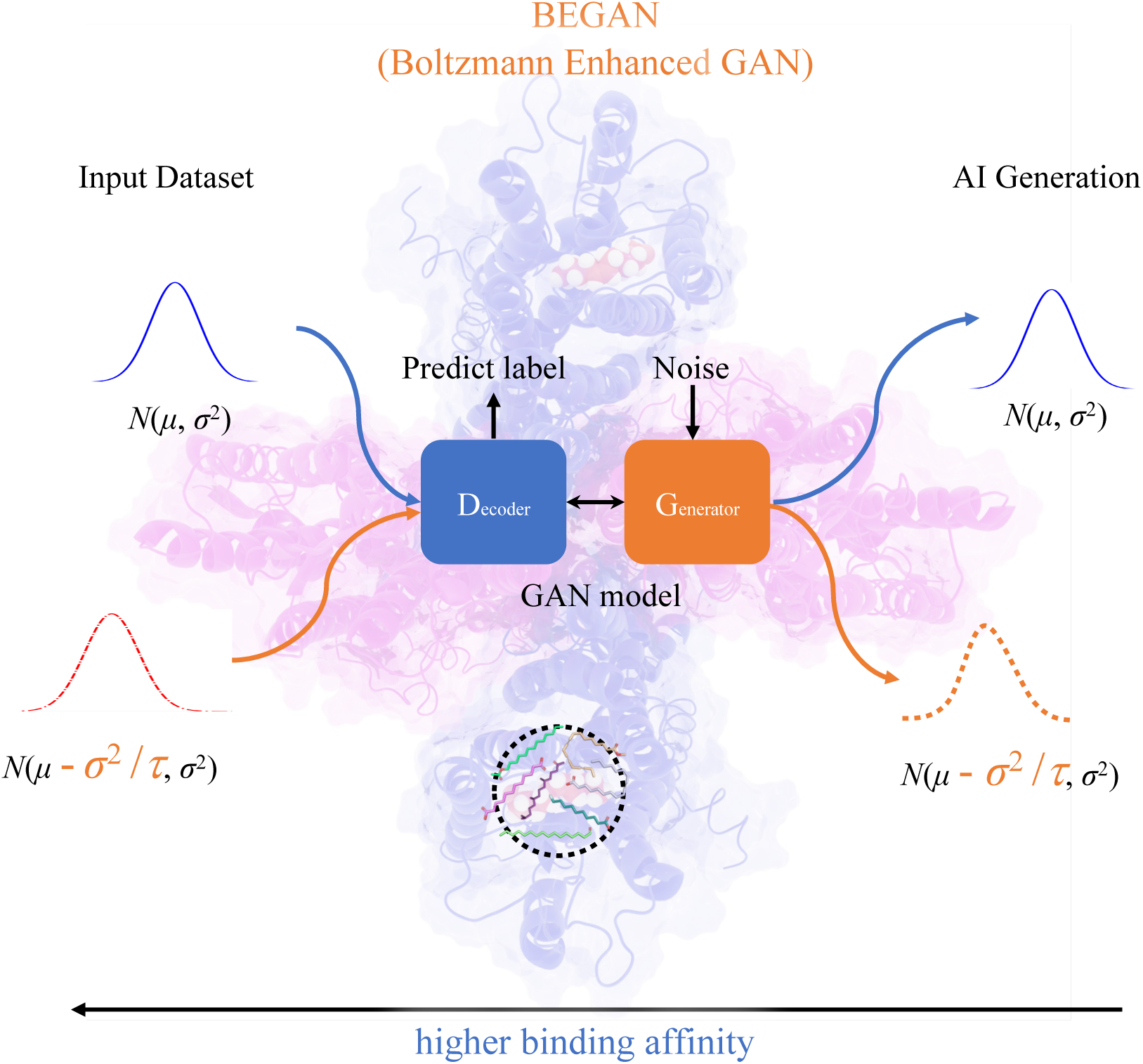

## Introduction

Drugs play a pivotal role in combating diseases, pests, and other challenges faced by humans over the long term. A crucial aspect of rational drug design involves identifying lead compounds that exhibit high binding affinity to target proteins. High-throughput screening methods, such as the docking technique,^1,2^ are effective for discovering these lead compounds within a restricted chemical space encompassing all potential molecules.

However, the challenge of defining such a confined chemical space persists due to the vast structural diversity of drug-like compounds, estimated at approximately 10^60^ possible structures.^3,4^ This difficulty has led to the establishment of extensive databases like PubChem^5^ and ZINC,^6^ which facilitate the identification of candidate molecules based on filter criteria such as molecular weight, similarity, lipophilicity (logP), and others. Nonetheless, the molecules currently cataloged in these databases represent only a small fraction of the vast chemical space.

An alternative approach involves constructing molecules by assembling atoms or molecular fragments guided by a fitness function using discrete local search methods like LUDI,^7^ MARS,^8^ and FEgrow.^9^ While effective, this method is limited in its ability to explore chemical space beyond the combinations present in initial libraries. ^10^

Artificial intelligence is now widely applied in drug design. Notable examples of this trend include machine learning models such as DeepFrag,^11^ DeepLinker,^12^ DeepScaffold,^13^ ScaffoldGraph,^14^ SurfGen,^4^ and Delete.^15^ These models utilize neural networks to generate molecules tailored to specific target proteins based on a local masked molecule, encompassing techniques like fragment elaboration, linker design, scaffold hopping, and side-chain decoration. They effectively identify candidates with high binding affinity. However, despite their success, these generative models seldom produce molecules with optimal binding affinity, and the distribution of binding affinities among the generated molecules remains stochastic. In most cases, the molecules generated by these models can undergo further optimization.

Generative Adversarial Networks (GAN) are well-known for their capability to align the generated data distribution (*p_gen_*) closely with the real data distribution (*p_data_*).^16^ However, traditional GAN, such as SeqGAN,^17^ often constrain the generated samples within the initial data distribution, making it challenging to produce molecules with high binding affinity that exist in smaller probability regions of the distribution. The objective-reinforced GAN (ORGAN) addresses this limitation by incorporating a weighted reward system where the generator optimizes a combination of domain-specific metrics and adversarial feedback from the discriminator.^18^ This approach fine-tunes distributions to generate molecules with improved drug-like properties and has also shown promise in material and drug discovery.^19^ In recent years, there has been considerable interest in investigating how insect odor receptors (ORs) recognize substrates,^20–25^ particularly in the context of developing targeted insect behavior regulators. ^26^ This research area has gained prominence with a focus on designing pheromone analog molecules, where artificial intelligence (AI) has proven to be a valuable tool. Insect ORs share a seven-transmembrane structure similar to mammalian G-protein-coupled receptors (GPCRs), albeit with an inverted orientation of the transmembrane helices—featuring a short intracellular N-terminus and an extracellular C-terminus. Research has primarily focused on insect ORs involved in detecting female-released sex pheromones, known as pheromone receptors (PRs), particularly in moths, which are known for their precise and sensitive sex pheromone communication systems. ^27^ Sex pheromones, volatile compounds detected by insect PRs like those found in moths, play a crucial role in guiding insects towards mating partners.^28^

Most characterized pheromone receptors (PRs) respond to Type I pheromones, which comprise 75% of natural pheromones. ^29–31^ These are primarily unsaturated C10–C18 fatty alcohols and their derivatives, such as acetates and aldehydes, with long, straight carbon chains. These compounds play a crucial role in attracting mates over short to moderate distances. Type II pheromones, accounting for 15% of natural pheromones, consist of longer, straight-chain C17–C25 unbranched polyunsaturated hydrocarbons and their epoxide derivatives.^22,29–32^ Unlike Type I pheromones, they lack functional groups but serve as long-range mate attractants. In addition, there are less common pheromones including Type III, which typically contain 1 or 2 methyl side-chain branches separated by an odd number of carbon atoms, and Type 0, which consists of short-chain alcohols or ketones with 7 to 9 carbon atoms.^31,33^ Designing sex pheromone analogs can significantly aid in moth capture and control by leveraging the regulation of insect olfactory behavior as an environmentally friendly pest management strategy.^22,26^

This study introduces BEGAN (Boltzmann-Enhanced GAN), a method for fine-tuning GANs to generate molecules with improved binding affinities. Using Boltzmann reweighting, the training data is adjusted based on estimated binding free energies obtained from a fast-docking algorithm, enhancing the likelihood of generating high-affinity molecules. The effectiveness of this approach is demonstrated through the customized design of pheromone analogs for the moth PR, *Spodoptera frugiperda* SfruOR16.

## Methods

### Training dataset preparation

We initially gathered 13 standard pheromones from iORbase^34^ and 36 previously identified pheromones^33^ (Table S1). To bolster the training of our neural network, we further enriched our dataset to include 13,308 molecules obtained through similarity searches in the PubChem database, ^5^ resulting in a total of 13,357 molecules in the training dataset (a full list is provided in the Supplementary Information).

**Table 1:** Metrics evaluating the quality of generated molecules across different reweighting iterations and scaling hyperparameter *τ*.

| | GAN | 1st Rew.<sup>a</sup> | 2nd Rew.<sup>b</sup> | 3rd Rew.<sup>c</sup> | $\tau = 0.3^d$ | $\tau = 0.2^e$ |
| --- | --- | --- | --- | --- | --- | --- |
| $P_{Val}$ | 93% | 94% | 95% | 95% | 91% | 94% |
| $P_{Uniq}$ | 90% | 86% | 78% | 60% | 69% | 31% |
| $P_{Nov}$ | 70% | 69% | 77% | 72% | 59% | 26% |
| $P_{V\&U}$ | 83% | 80% | 73% | 55% | 60% | 26% |
| $P_{V\&U\&N}$ | 64% | 61% | 62% | 50% | 47% | 19% |

The superscripts a, b, and c denote the first, second, and third rounds of reweighting with $\tau = 0.6$ , respectively. The superscripts d and e indicate the first round of reweighting with $\tau = 0.3$ and $\tau = 0.2$ , respectively.

### Enhance the training data by Boltzmann reweighting

The general outline of our method, BEGAN (Boltzmann-Enhanced GAN), is depicted in Figure 1. BEGAN serves as a preprocessing step for the GAN-based molecular generation model, which aims to generate a distribution across the molecular space consistent with the input dataset. The probability of a molecule binding to a protein is determined by its binding free energy. According to the Boltzmann distribution, this probability is exponentially related to the negative of the binding free energy, scaled by a temperature-dependent constant: exp(*−*Δ*U/τ*), where Δ*U* represents the estimated binding free energy and *τ* is the temperature-related scaling factor, which for simplicity is represented as *k*_B_*T* in units of kcal/mol. Consequently, the relative binding probabilities of two molecules with binding free energies Δ*U*_1_ and Δ*U*_2_ are proportional to their respective exponential terms, as shown in Figure 1(a).

**Figure 1:**
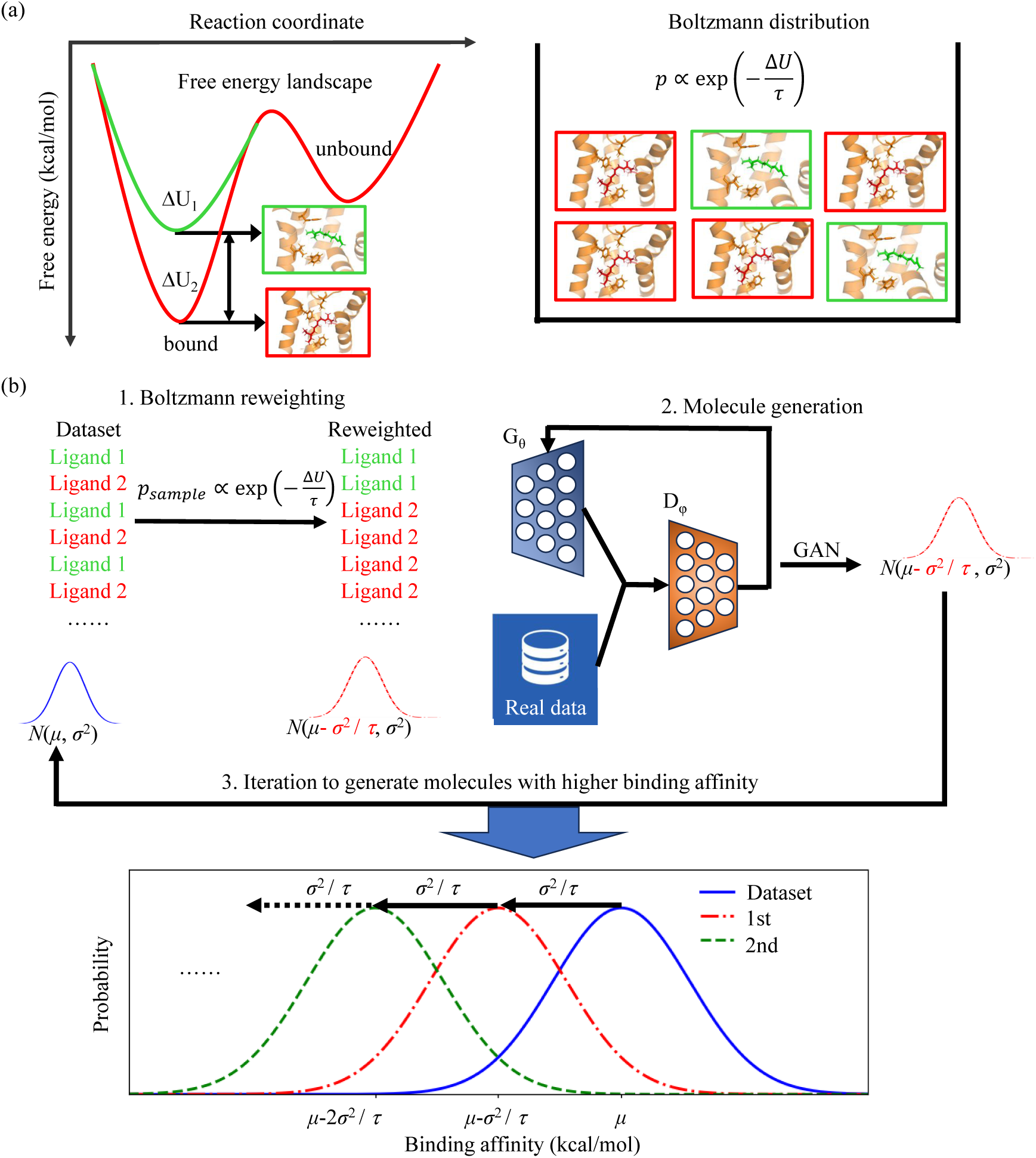
Schematic overview of dataset augmentation via Boltzmann reweighting. (a) Free energy landscape illustrating the relationship between binding affinity and ligand population. Ligand 1 with higher free energy (Δ*U*_1_) has a smaller population compared to ligand 2 with smaller free energy (Δ*U*_2_). The relative abundance of ligands is determined by the Boltzmann factor (*e^−^*^Δ^*^U/τ^*), where *τ* is a scaling hyperparameter related to temperature. (b) Pipeline for generating molecules with enhanced binding affinities: 1) Boltzmann Reweighting: Adjusts the distribution of ligands in the original dataset based on their binding affinities using the Boltzmann factor. 2) Molecule Generation: Employs a GAN-based model to generate new molecules. 3) Iteration: Repeatedly applies reweighting and generation steps to progressively shift the generated molecule distribution towards higher binding affinity regions.

To increase the likelihood of generating molecules with stronger binding affinities during training, we adjusted the training data weights based on their binding free energies, with the probability *p_sample_* ∝; exp(*−*Δ*U/τ*). This reweighting assigns higher probabilities to molecules with smaller Δ*U*, indicating higher binding affinity. By using this adjusted dataset as input to the neural network, we aim to generate a batch of molecules that reflects the binding affinity distribution observed in the reweighted input dataset.

We further demonstrate that the reweighting process primarily shifts the mean of the distribution without altering its shape or variance (as discussed in Equation 2 and related sections), which allows us to iterate the reweighting process. By reweighting the generated molecules and using them as inputs for subsequent iterations, we can progressively refine the distribution, steering it towards higher binding affinities. This iterative process, depicted in Figure 1(b), enables the identification of molecules with increasingly improved binding affinities over successive iterations.

### Molecular generation model

To prepare molecules for recognition by neural networks, we employed SMILES encoding^35,36^ to represent them as character strings that can be easily hot-encoded into binary matrices. It is important to note that SMILES representations are non-unique, meaning several strings may represent the same molecule. Therefore, adopting a standardized SMILES format such as kekuleSmiles^35^ or canonical SMILES,^36^ which are well-integrated into the RDKit^37^ package, is preferable. GANs^16^ are effective for generating molecular structures that mimic a specified initial distribution. They are a typical generative model designed to minimize the Jensen-Shannon (JS) divergence between the real data distribution *p_data_* and the generated distribution *p_gen_* produced by the generator *G_θ_* (eq. 1).^38^ When the JS divergence approaches 0, the distribution *p_gen_* converges to *p_data_*.

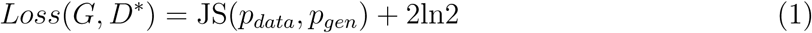

In this work, we employ the ORGAN model as the molecular generation tool, maintaining drug-likeness while focusing on further optimizing the binding affinity of generated molecules using our Boltzmann reweighting protocol.

### Structural prediction and molecular docking

The exact stoichiometry of OR and Orco complexes remains a topic of ongoing debate, with recent cryo-EM studies suggesting predominant heterotetrameric assemblies of 1OR:3Orco and potentially 2OR:2Orco. ^23,24^ In this study, we employed a 2OR:2Orco (as shown in Figure 4(a)) assembly arranged in a diagonal opposite conformation, consistent with our previous methodology.^21,22^

The structure of SfruOR16 (UniProt ID: A0A9R0DR18) was predicted using AlphaFold v2.3.2.^39^ Utilizing this predicted heterotetrameric structure of the SfruOR16-SfruOrco complex, we modeled the ligand-bound complex by docking ligands with the SfruOR16 monomer using AutoDock Vina-GPU.^2^ The binding pocket, identified using fpocket,^40^ was consistent with our previous models of HarmOR14b and HarmOR16, ^21^ serving as the docking site for subsequent docking. The grid dimensions were set to (18 Å, 18 Å, 18 Å).

### Molecular dynamics simulations of ligand-bound OR/Orco heterotetramer

To further validate the model, molecular dynamics (MD) simulations were performed on the two top-ranked pheromone analogs molecules in complex with moth pheromone receptors (PRs), as depicted in Figure 2(d). The conformations of the top 100 ranked pheromone analogs molecules are presented in Figure S1 with their corresponding SMILES strings provided in Table S1. Each OR/Orco heterotetramer accommodates two ligands, enabling the generation of two ligand-binding trajectories from a single simulation of the complex.

**Figure 2:**
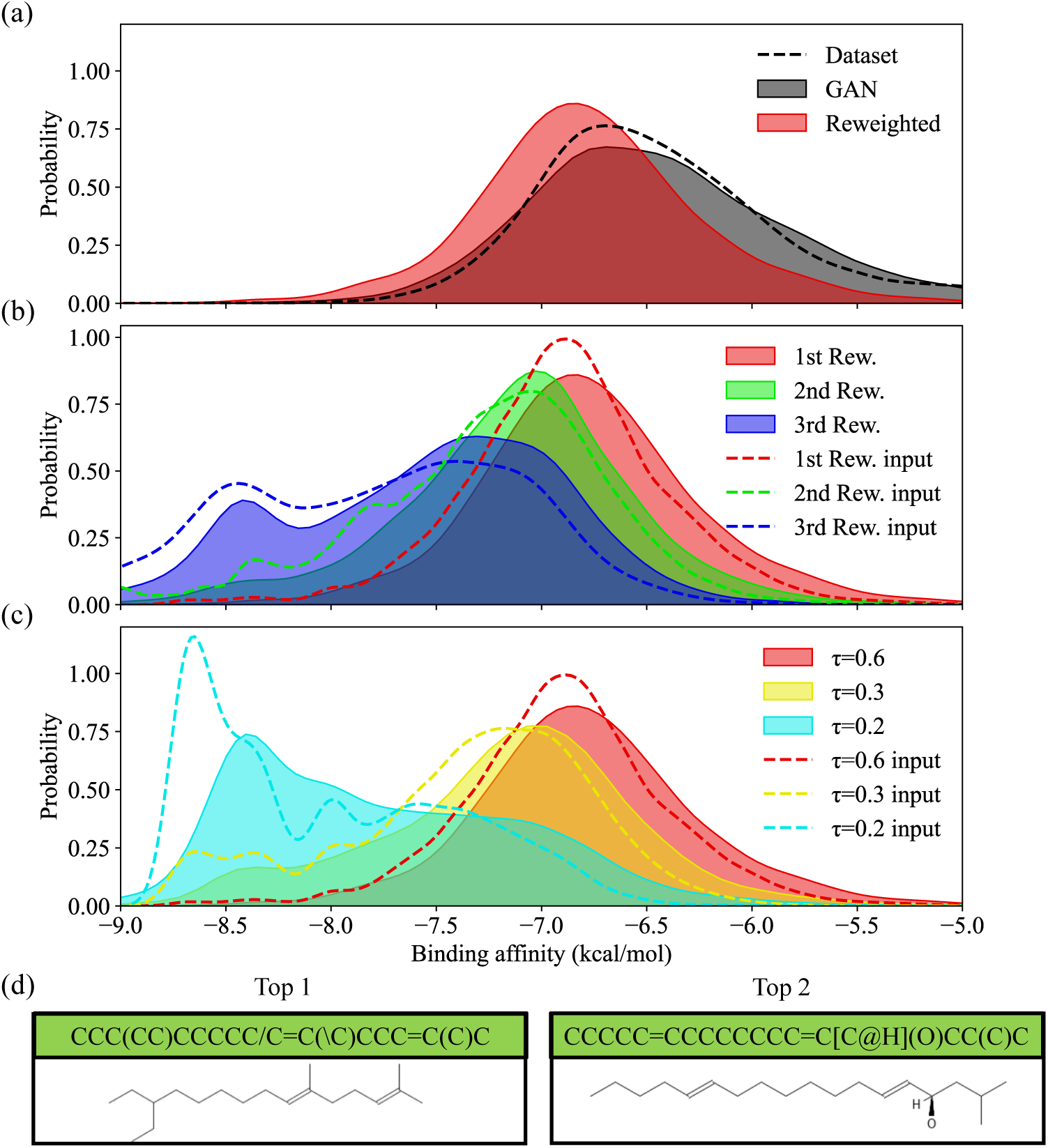
Dataset augmentation for pheromone analogs molecule generation via Boltzmann reweighting. (a) Binding affinity distribution of molecules in the initial dataset (black dotted line), generated by GAN (black area), and generated by GAN after one round of reweighting at *τ* = 0.6 (red area). (b) Binding affinity distribution of molecules in the training set after reweighting once (red), twice (green), and three times (blue) at *τ* = 0.6 (dotted lines), and generated by GAN after reweighting once (red), twice (green), and three times (blue) at *τ* = 0.6 (shaded areas). (c) Binding affinity distribution of molecules in the training set after reweighting at *τ* = 0.6 (red), 0.3 (yellow), and 0.2 (cyan) (dotted lines), and generated by GAN after one round of reweighting at *τ* = 0.6 (red), 0.3 (yellow), and 0.2 (cyan) (shaded areas). (d) Two representative pheromone analogs from the top 100 molecules generated by GAN after three rounds of reweighting and pheromone feature-based filtering (see Figure S1 for details).

The insertion of protein-ligand complexes into a lipid bilayer composed of 1-palmitoyl-2-oleoyl-sn-glycero-3-phosphocholine (POPC), solvated with TIP3P water and 0.15 M NaCl, was facilitated using CHARMM-GUI.^41,42^ The protein was modeled using the CHARMM36m force field,^43^ and parameters for the ligands were generated using the CHARMM General Force Field (CGenFF).^44^ The initial conformation underwent an energy minimization step followed by six equilibration steps to achieve relaxation. Temperature control at 310 K with a 1 ps coupling constant was applied using the velocity-rescale thermostat, and pressure was maintained at 1 bar with a 5 ps time coupling constant using Parrinello-Rahman pressure coupling scheme. Van der Waals interactions were cut off at 1.2 nm with a switch function starting at 1.0 nm, while short-range electrostatic interactions were handled similarly. Long-range electrostatics were computed using the particle mesh Ewald (PME) algorithm with a mesh spacing of 0.12 nm. For production simulation, each system underwent two independent 500 ns semi-isotropic pressure coupling simulations. Since each complex consists of two identical ligand-bound subunits, the total simulation time for each model amounted to 2 *µ*s.

All simulations were conducted using a GPU-accelerated version of Gromacs 2022.5. ^45^ The interaction between proteins and ligands was analyzed using GetContacts (https://getcontacts.github.io/), and the root-mean-square deviation (RMSD) of the ligands was calculated using the Gromacs gmx tool.

### Binding affinity calculated by MM/PBSA

The energetic contribution of residues within 5 of the ligand was estimated using Molecular Mechanics / Poisson Boltzmann (Generalized Born) surface area (MM/PB(GB)SA) methods,^46,47^ following the per-residue effective free energy decomposition protocol. For each complex, the last 50 ns of the MD trajectories were used for MM/PBSA calculations. The total molecular mechanics energy, Δ*E*_MM_, includes bonded, electrostatic, and van der Waals interactions. The electrostatic solvation energy was calculated using the Poisson Boltzmann model (Δ*G*_PB_), and the non-polar contribution was estimated based on a linear relation with the solvent-accessible surface area (Δ*G*_SA_). The residue-specific energetic contributions were obtained by averaging the energy decomposition analysis results from the two independent trajectories.

## Result

### Dataset Preparation and Initial GAN Training

We used the moth pheromone receptor SfruOR16 from the fall armyworm *Spodoptera frugiperda* as the model system to test our optimized molecule generation method. To expand our neural network training dataset, we first collected a total of 13,357 molecules with pheromone-like properties, primarily sourced from the PubChem database through similarity searches,^5^ supplemented by additional identified pheromones from iORbase^34^ and previous reports.^33^ Following the augmentation of our dataset, we evaluated the binding affinities of all molecules using the AutoDock Vina-GPU molecular docking algorithm.

The initial distribution of binding affinities for these molecules in the original training dataset is depicted in Figure 2(a) (black dashed line). These molecules served as the initial training set for the GAN, from which 10,000 molecules were randomly selected post-training for docking to estimate their binding affinities. Note that we always used 10,000 molecules from the generated dataset for subsequent docking if without specification. The binding affinity distribution of molecules generated by the GAN is shown in Figure 2(a) (grey area). The distribution of the input dataset and the GAN-generated dataset are nearly identical, indicating successful minimization of the Jensen-Shannon (JS) divergence between the distributions *p_data_* and *p_gen_* (as described in Equation 1).

### Enhance the training dataset by Boltzmann reweighting

To enhance the likelihood of generating molecules with stronger binding affinities during training, we adjusted the weights of the training data based on their estimated binding free energies (Δ*U*), favoring molecules with smaller Δ*U* for stronger binding. Utilizing the Boltzmann factor, exp(*−*Δ*U/k*_B_*T*), where k_B_ is the Boltzmann constant and *T* is temperature, we recalibrated the occurrence probabilities with *p_sample_* ∝ exp(−Δ*U/τ*) at *τ* = 0.6, as illustrated in Figure 1(b). This enhanced dataset was subsequently used to train a GAN. Additionally, since the reweighting process does not alter the shape or variance of the distribution, it can be further refined by iterating the reweighting process, as demonstrated in Equation 2. As shown in Figure 2(a), the resulting molecules exhibited a significantly improved binding affinity distribution compared to the original dataset, demonstrating the effectiveness of our weighting strategy.

We iteratively reweighted the generated molecules twice more using *τ* = 0.6, training a GAN on each resulting dataset. The binding affinity distributions of these doubly and triply reweighted datasets are shown in Figure 2(b) as green and blue areas, respectively. Our results indicate that successive reweighting effectively shifted the distribution towards higher binding affinities. Nevertheless, the emergence of a second peak in the high-affinity region after the third iteration suggested potential overfitting due to limited high-affinity data in the original dataset. Therefore, we terminated the iterative process at this point.

### Effect of Hyperparameter ***τ*** on GAN Performance

The hyperparameter *τ* significantly influences how much the generated molecule distribution diverges from the original dataset. To investigate this, we examined the binding affinity distributions of molecules generated by the GAN after one round of reweighting at *τ* values of 0.6, 0.3, and 0.2, as visualized in Figure 2(c). Our analysis revealed that decreasing *τ* generally improves the GAN’s ability to generate high-affinity molecules. However, excessively small *τ* values, such as 0.2 (yellow area in Figure 2(c)), can lead to overfitting, where the generated distribution (shadowed area in Figure 2(c)) deviates considerably from the original data distribution (dotted lines).

Figure 2(d) showcases the two top-ranked molecules with desirable pheromone analog properties generated by the GAN after three reweighting iterations.

### Evaluating the Impact of Reweighting on Molecule Generation Quality

GANs can sometimes produce redundant or invalid molecules due to limitations in training data coverage, which might not encompass all valid chemical structures. This can lead to the generation of repetitive or chemically unsound molecules. To evaluate the impact of the hyperparameter *τ* on the quality of generated molecules, we introduced metrics for novelty, uniqueness, and validity:

- *P*_Val_: The percentage of generated molecules that are valid according to specified criteria, such as syntactic correctness in SMILES strings or adherence to chemical rules.
- *P*_Uniq_: The percentage of generated molecules that are unique and non-redundant. This metric measures the diversity of the generated molecules.
- *P*_Nov_: The percentage of generated molecules that do not exist in the initial training dataset. This indicates the model’s ability to create novel molecules.
- *P*_V&U_: The percentage of generated molecules that are both valid and unique, combining validity and uniqueness to assess the model’s ability to produce chemically sound and diverse molecules.
- *P*_V&U&N_: The percentage of generated molecules that are valid, unique, and not present in the initial training dataset. This metric represents the model’s capability to generate novel, chemically valid molecules distinct from known molecules.

Our analysis, detailed in Table 1, indicates that the Boltzmann reweighting process has minimal impact on the validity ratio, suggesting that the GAN consistently generates valid SMILES representations. However, at a small scaling hyperparameter of *τ* = 0.2, only 26% of the generated molecules are novel, with most being present in the training set, suggesting overfitting. When considering both validity and uniqueness, this ratio decreases to 19%. The metric *P*_V&U&N_ remains consistently high after the first and second rounds of reweightings, though it is slightly lower than the GAN baseline. In the third round, as the distribution shifts further towards the high-affinity region, *P*_V&U&N_ drops significantly to around 50%.

We further checked the binding affinity of the generated molecules to SfruOR16, as shown in Figure 3(a). The results suggested that a small *τ* (*τ* = 0.2) struggles to generate a sufficient number of unique, high-affinity molecules, leading to predominantly redundant molecules. While the mean binding affinity at *τ* = 0.2 is comparable to that at *τ* = 0.3, the number of molecules outperforming the initial dataset is minimal (Figure 3(b, c)). In contrast, iterative reweighting consistently improves mean binding affinity and the count of molecules surpassing the initial dataset’s best performer. This approach enables the GAN to identify and generate novel, high-affinity molecules. Our findings suggest that moderate *τ* values (e.g., 0.6) provide a balance between exploration and exploitation, allowing the GAN to generate diverse molecules while maintaining focus on high-affinity regions. Smaller *τ* values limit the model’s exploration capabilities.

**Figure 3:**
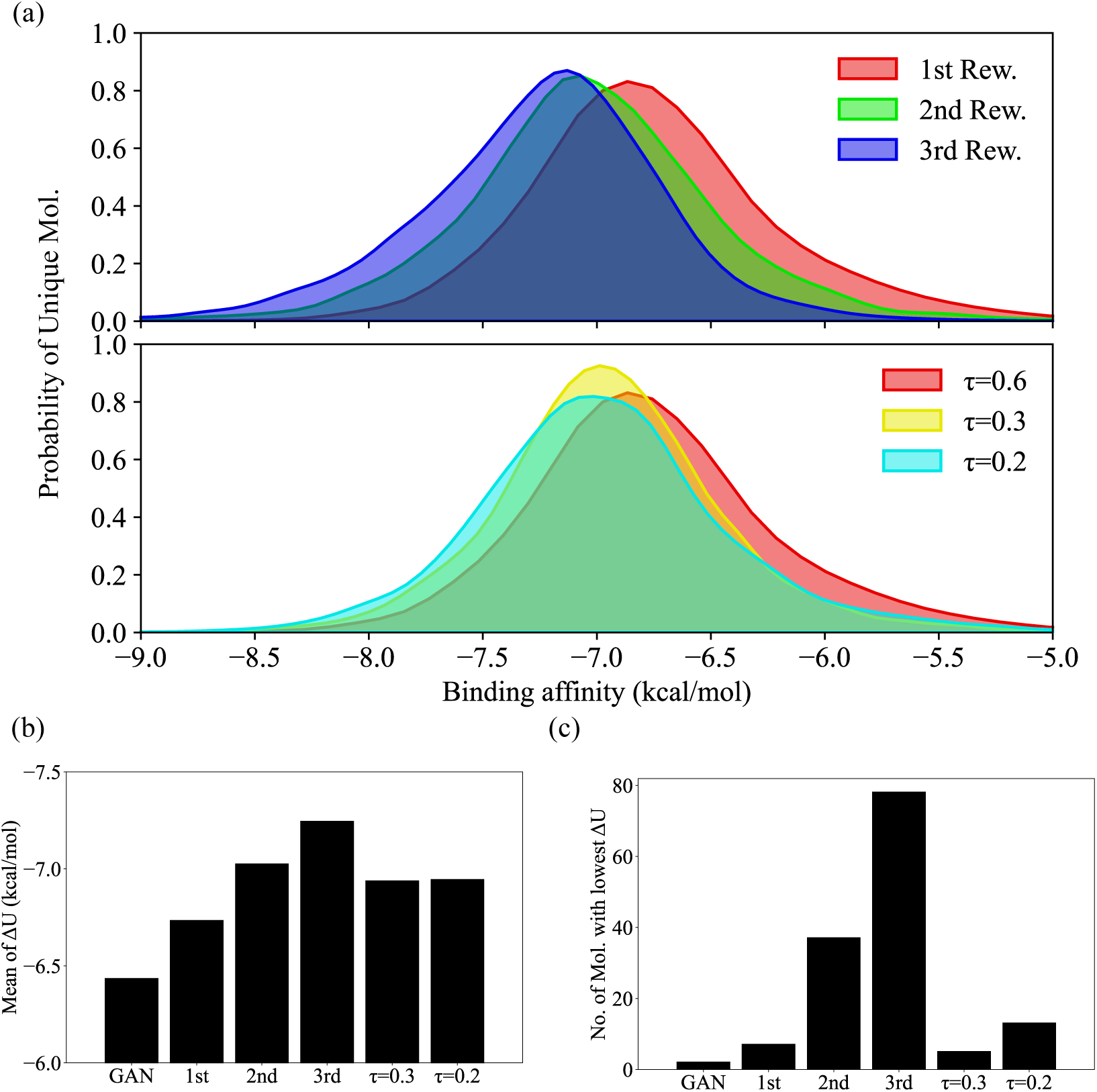
The effect of hyperparameter τ on the molecule generation. (a) Upper panel: Binding affinity distributions after one (red), two (green), or three (blue) rounds of reweighting at *τ* = 0.6. Lower panel: Binding affinity distributions after one round of reweighting at *τ* = 0.6 (red), 0.3 (yellow), or 0.2 (cyan). In both panels, distributions are represented by solid lines with shaded areas. (b) Mean binding affinities of 10,000 non-repeating molecules across different conditions; (c) Number of molecules with binding affinities exceeding the initial dataset’s best (−8.7 kcal/mol) among 10,000 unique molecules, indicating the GAN’s exploration of chemical space relative to the original dataset.

Additional analyses using different Δ*U* cutoffs (Figure S2) support these conclusions. Iterative reweighting at *τ* = 0.6 consistently outperforms other conditions in generating unique, high-affinity molecules. This emphasizes the importance of iterative optimization and parameter tuning for effective molecule generation and drug discovery.

### Binding Dynamics and Stability Analysis using MD Simulations

To assess the dynamic binding behavior of the generated molecules, we performed molecular dynamics (MD) simulations on the pheromone-bound SfruOR16/Orco complex. The top two molecules from the third reweighting iteration (Figure 2(d), named Top1 and Top2) were selected and compared to the experimentally validated pheromone Z9-14:OH (IUPAC name:(Z)-tetradec-9-en-1-ol; CID: 5364712). Previous studies have revealed that SfruOR16 exhibits a strong response to Z9-14:OH. ^48^

MD simulations confirmed that all ligands remained stably bound within the receptor pocket (Figure 4(a)). Notably, Top1 and Top2 exhibited smaller root mean square deviations (RMSDs) than Z9-14:OH (Figure 4(b) and Figure S3), indicating greater conformational stability for the AI-designed molecules. In addition, the binding affinities estimated by MM/PBSA (Figure 4(c)) for the designed molecules were higher than those of Z9-14:OH, supporting that both Top1 and Top2 molecules might have enhanced binding affinity and stability compared to the experimentally identified pheromones.

**Figure 4:**
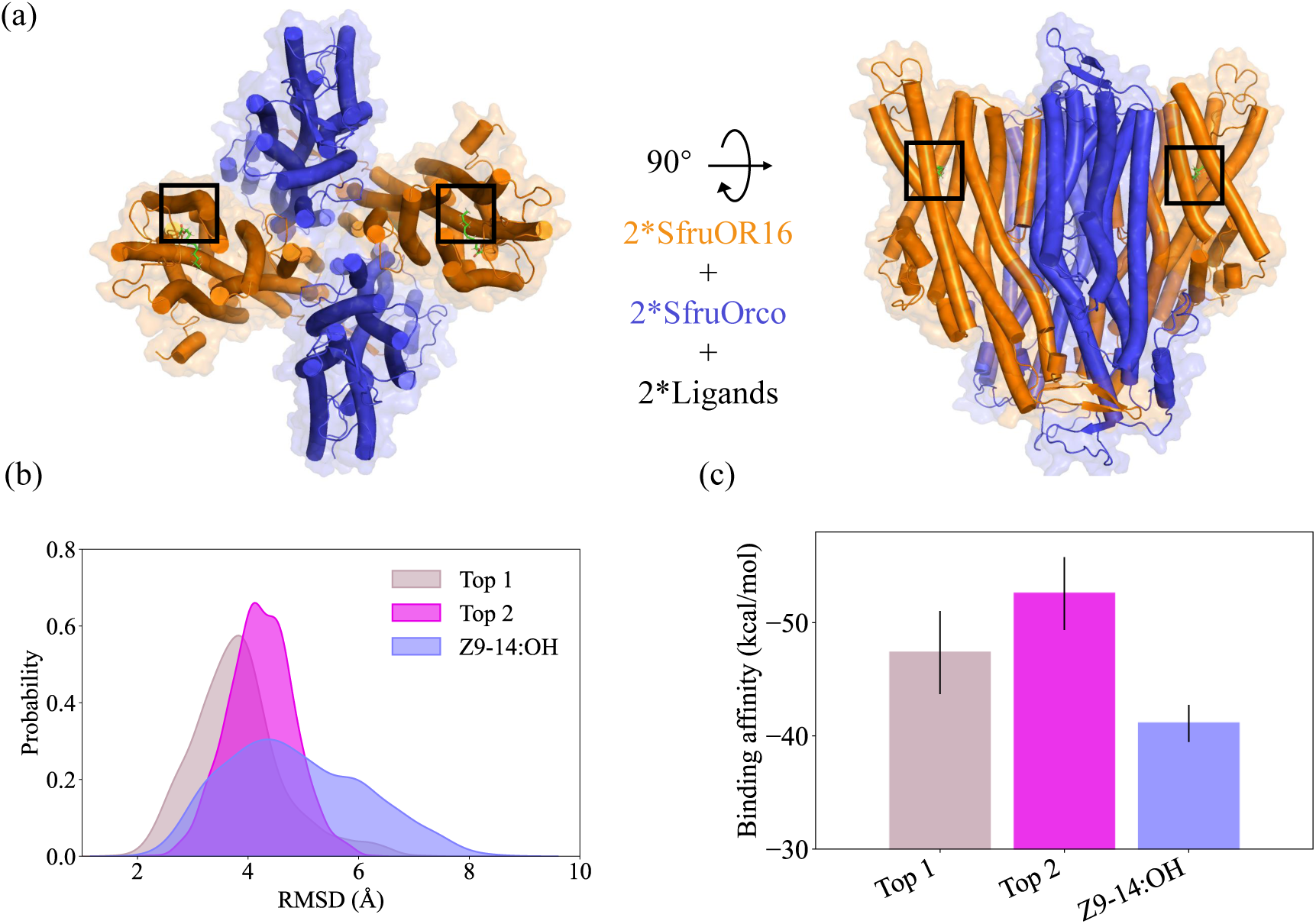
Molecular dynamics simulations of SfruOR16/Orco heterotetramer-ligand complexes. (a) Top view of the OR/Orco heterotetramer (left) and side view of the OR/Orco heterotetramer (right) with the binding site highlighted by a black box. (b) Ligands RMSD distribution derived from four 500 ns MD trajectories. Note that the ligand RMSD was calculated after aligning the MD conformations with the heterotetramer backbone. (c) Binding affinities of these ligands estimated by MM/PBSA.

The MD simulations further demonstrated that all ligands maintained a stable state within the binding pocket throughout the two repeats of MD simulation (Figure S4(a-c)). Protein-ligand interaction analysis revealed that all the molecules primarily engage in hydrophobic interactions with the receptor (Figure S4(d-f)). Notably, both Top2 and Z9-14:OH formed hydrogen bonds with SfruOR16, with the designed molecules forming these hydrogen bonds slightly more frequently than Z9-14:OH, which may contribute to their enhanced binding affinity (Figure S4(e,f)).

These findings collectively demonstrate the superior binding properties of the GAN-generated molecules, highlighting the potential of our approach for designing potent pheromone analogs.

## Discussion

### Theoretical Framework for Iterative Reweighting

The core principle of GAN is to generate data that closely resembles the input dataset. Our approach focused on shifting the distribution of generated molecules *p_gen_* towards higher binding affinity regions by reweighting the training data. This involved assigning probabilities to molecules based on their binding affinities using the Boltzmann factor, *p_sample_* ∝ exp(*−*Δ*U/τ*), where Δ*U* is the binding free energy and *τ* is a temperature-like scaling parameter, which will be elaborated on later.

We observed that the binding affinity distribution approximated a normal distribution (Figure 2(a)). Assuming the initial dataset follows a normal distribution *p_data_ ~ N* (*µ, σ*^2^), theoretical analysis (as shown in Equation 2) shows that reweighting shifts the distribution *p_re_* to *N* (*µ − σ*^2^*/τ, σ*^2^), with the mean decreasing by *σ*^2^*/τ*. Consequently, GAN is expected to generate molecules following a distribution centered around *N* (*µ − σ*^2^*/τ, σ*^2^). Subsequent reweighting further shifts the distribution by 2*σ*^2^*/τ*.

This iterative process allows for the progressive enhancement of the dataset towards higher binding affinities. By carefully adjusting *τ* and applying iterative reweighting, we can effectively control and optimize the distribution of generated molecules to match desired properties. This approach not only improves molecule quality but also demonstrates GAN’s adaptability for iterative dataset refinement.

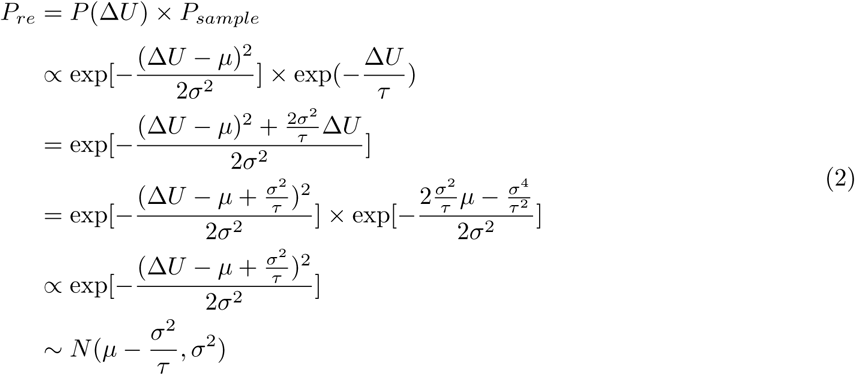

### Optimal Hyperparameter Selection of ***τ*** for Iterative Reweighting

The hyperparameter *τ* plays a crucial role in determining the extent to which the distribution is shifted. A practical choice for *τ* is 0.6, corresponding to the value of thermal energy at room temperature. However, *τ* can be adjusted based on specific needs. Smaller *τ* values result in sharper distributions, while higher values lead to more uniform distributions.

Reweighting shifts the distribution leftward by *σ*^2^*/τ*. For *τ* values of *σ*, *σ/*2, and *σ/*3, the shift is *σ*, 2*σ*, and 3*σ*, respectively. Considering the 3*σ* rule of statistics, shifting the distribution by 3*σ* is impractical as it would primarily generate outliers.

In the context of pheromone analogs design, with *σ ≈* 0.6, *τ* values of 0.6, 0.3, and 0.2 correspond to shifts of *σ*, 2*σ*, and 3*σ*. A *τ* of 0.2 led to severe overfitting (Figure 3(a)) due to the extremely small probability of observing molecules beyond 3*σ* in the training data. This limited the GAN’s ability to explore novel chemical space. A *τ* of 0.3 showed slight overfitting (Figure 3(a)), while 0.6 exhibited minimal overfitting (Figure 2(b)). Excessive iterative reweighting can also lead to overfitting, as evidenced by the stagnation of novelty values (Table 1).

Throughout the iterative process, maintaining alignment between the input and output data distributions, as depicted in Figure 2(b), reinforces the efficacy of choosing *τ* = 0.6 or *σ*, based on our findings. While *τ* = *σ/*2 is feasible but not recommended.

## Conclusion

In this study, we have developed a method called BEGAN (Boltzmann-Enhanced GAN) to enhance Generative Adversarial Networks (GAN) in generating molecules with high binding affinity. Leveraging recent advancements in GAN, which aim to bridge the gap between input data distribution *p_data_* and generated distribution *p_gen_*, we employed a reweighting strategy to shift the generated distribution *p_gen_* towards molecular spaces with higher binding affinity. This approach, rooted in Boltzmann distribution principles, adjusted the occurrence frequency of molecules during training according to *p_sample_* ∝ exp(*−*Δ*U/τ*), where Δ*U* represents the binding free energy of a molecule to its target protein, and *τ* controls the magnitude of the distribution shift. Theoretically, iterative reweighting can progressively refine the distribution *p_gen_*, potentially uncovering molecules with higher binding affinities within the dataset. Our analysis of different *τ* values highlights the importance of careful parameter selection for optimal performance. We recommend using *τ* values of 0.6 or *σ* (standard deviation of the initial data distribution) for most applications.

Through iterative refinement, we systematically identified novel molecules with the highest binding affinities. Moreover, the top two molecules designed by the GAN after three rounds of reweighting showed improved binding affinities with SfruOR16 compared to the experimentally identified pheromones Z9-14:OH. Atomistic MD simulations further validated that our reweighting strategy effectively enhanced the GAN’s capability to generate both sex pheromone analogs with superior binding affinities to SfruOR16. These findings suggest their potential as potent sex pheromone analogs, pending further experimental validation.

This work demonstrates the potential of our reweighting strategy for discovering novel drug candidates. As docking methodologies continue to advance, we anticipate that our BEGAN approach will remain effective in identifying high-affinity molecules across various target proteins.

## Code availability

The source code of the Boltzmann reweighting and MD input files are available at GitHub (https://github.com/yongwangCPH/papers/tree/main/2024/BoltzmannReweightingDrugDesign)

## Acknowledgements

We acknowledge the financial support of the National Natural Science Foundation of China (No. 32371300 to Y.W., 12174337 and 11932017 to X.P.), the Zhejiang Provincial National Science Foundation of China (No. LZ24C050003 to Y.W.), and the National Key Research and Development Program of China (No. 2021YFF1200404 to Y.W.).

## Supporting Information

The following files are available free of charge.

- initial_dataset.xlsx: The molecules in the initial dataset collected from iORbase, previous reports, and similarity searches in PubChem.
- 1th.xlsx: The SMILES strings of the molecules generated by the GAN after one round of reweighting at *τ* = 0.6.
- 2th.xlsx: The SMILES strings of the molecules generated by the GAN after two rounds of reweighting at *τ* = 0.6.
- 3th.xlsx: The SMILES strings of the molecules generated by the GAN after three rounds of reweighting at *τ* = 0.6.
- T=0.2.xlsx: The SMILES strings of the molecules generated by the GAN after one round of reweighting at *τ* = 0.2.
- T=0.3.xlsx: The SMILES strings of the molecules generated by the GAN after one round of reweighting at *τ* = 0.3.

**Figure S1:**
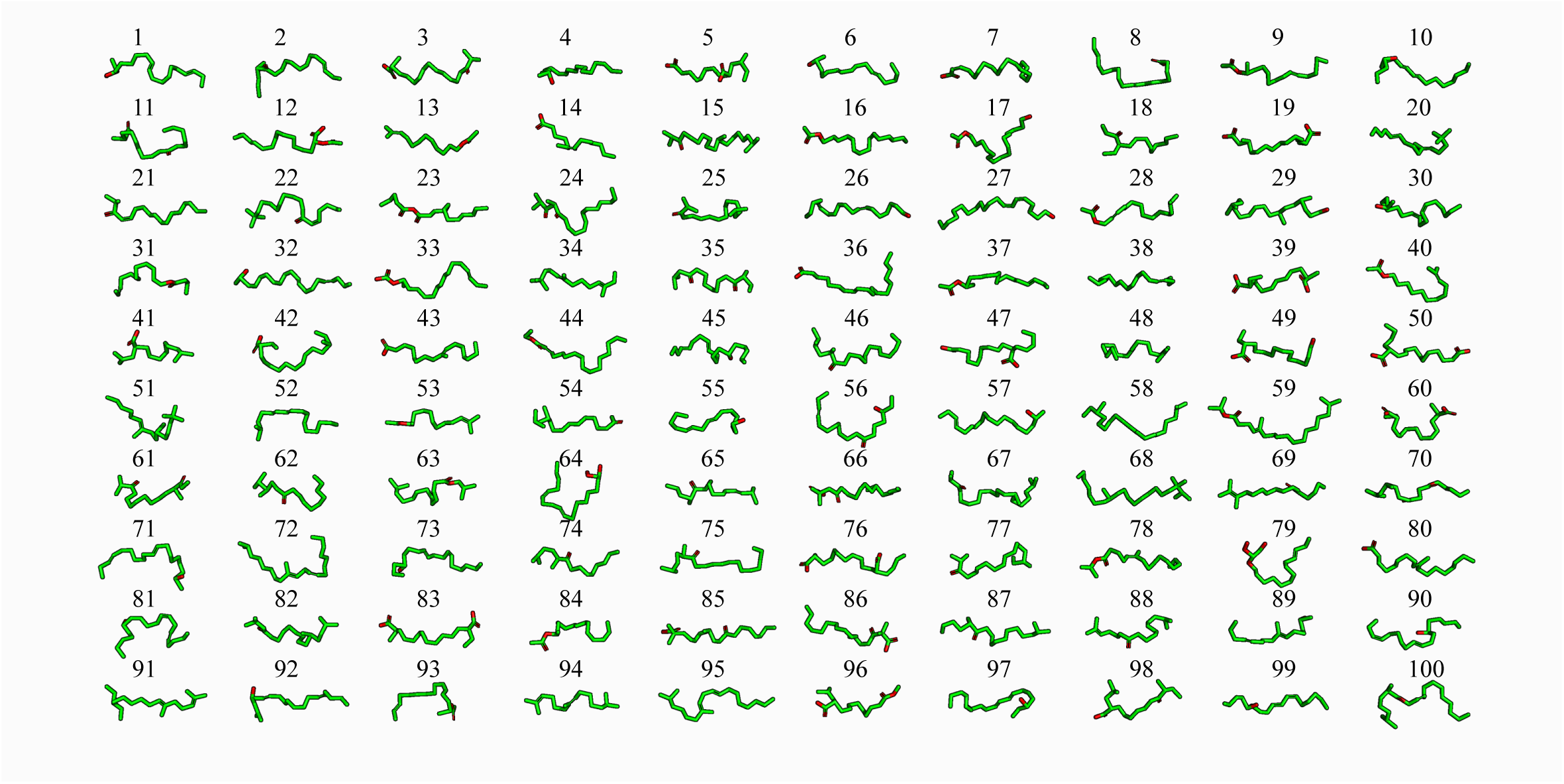
Illustrations of the top 100 AI-generated pheromone analogs and its analog molecules. The corresponding SMILES for these molecules can be found in Table S2. The molecules were generated via a GAN-based approach involving three rounds of reweighting.

**Figure S2:**
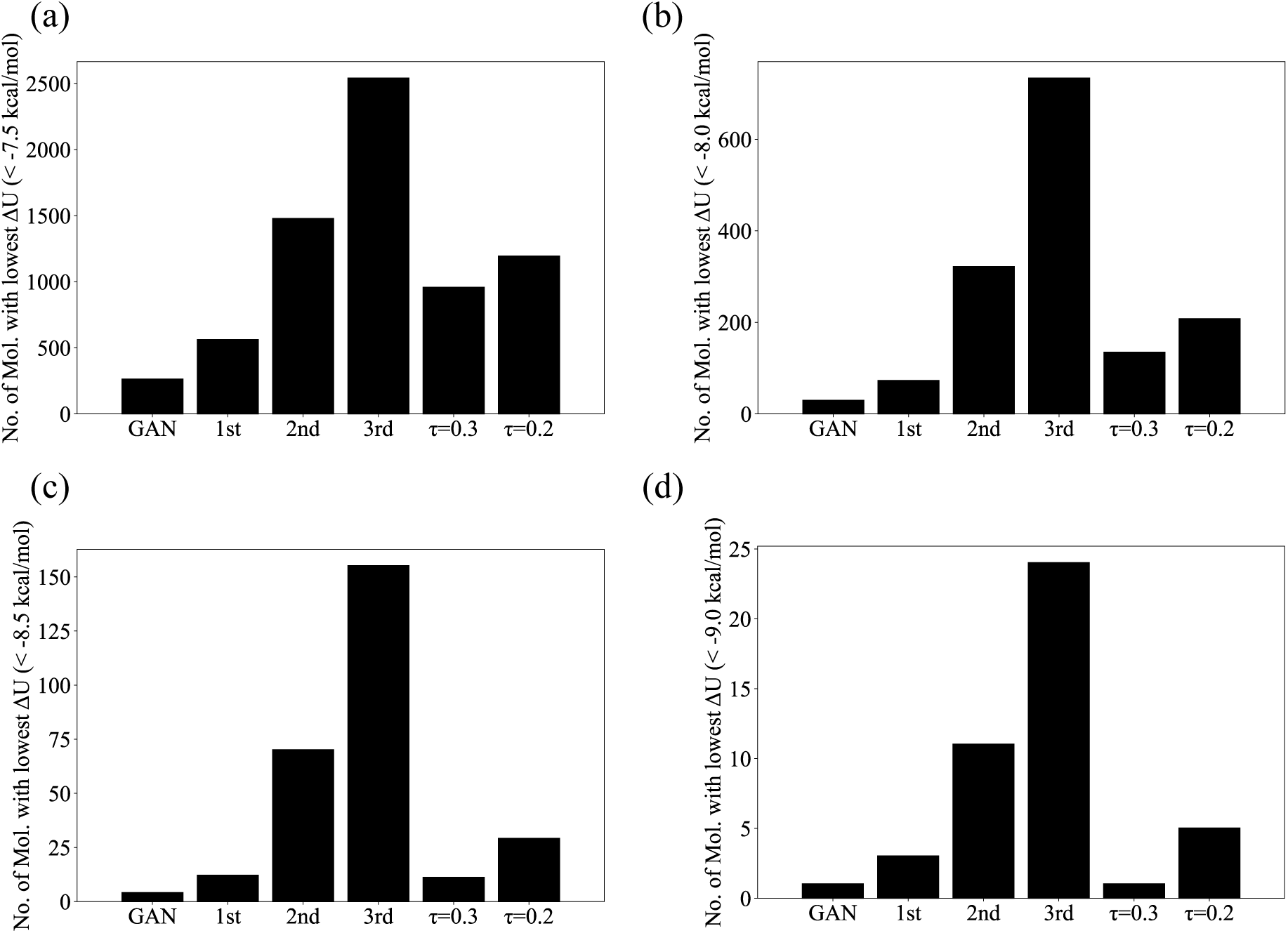
Number of molecules with binding affinities below various thresholds among 10,000 unique molecules. Panels (a-d) show the number of molecules exhibiting binding affinities smaller than −7.5, −8.0, −8.5, and −9.0 kcal/mol, respectively.

**Figure S3:**
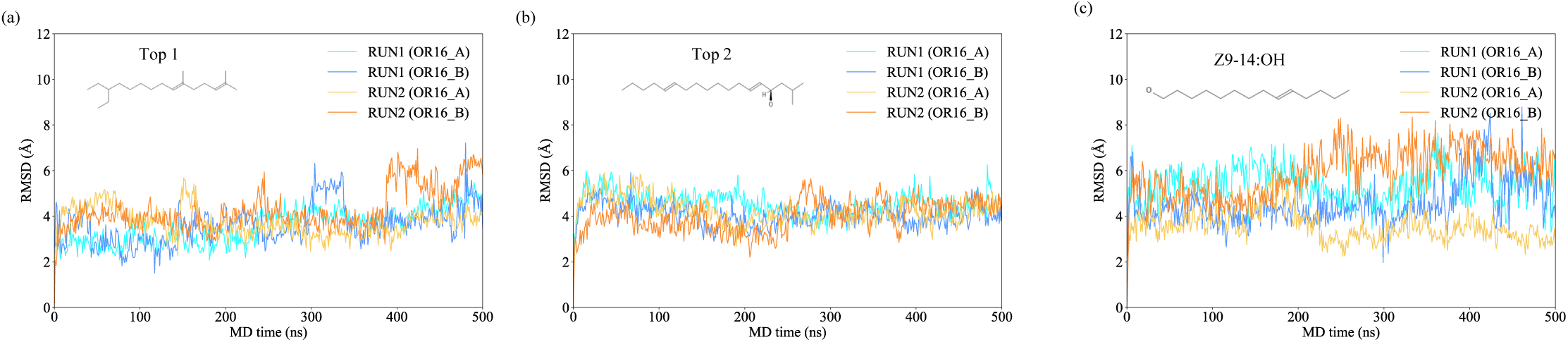
Trajectories of root mean square deviation (RMSD) of ligands from their initial conformation during MD simulations of SfruOR16-Orco heterotetramers. Panels (a-c) display RMSD trajectories for Top 1,2 and Z9-14:OH ligands. MD conformations were aligned to the protein structure of the heterotetramer backbone before RMSD calculation of the ligands.

**Figure S4:**
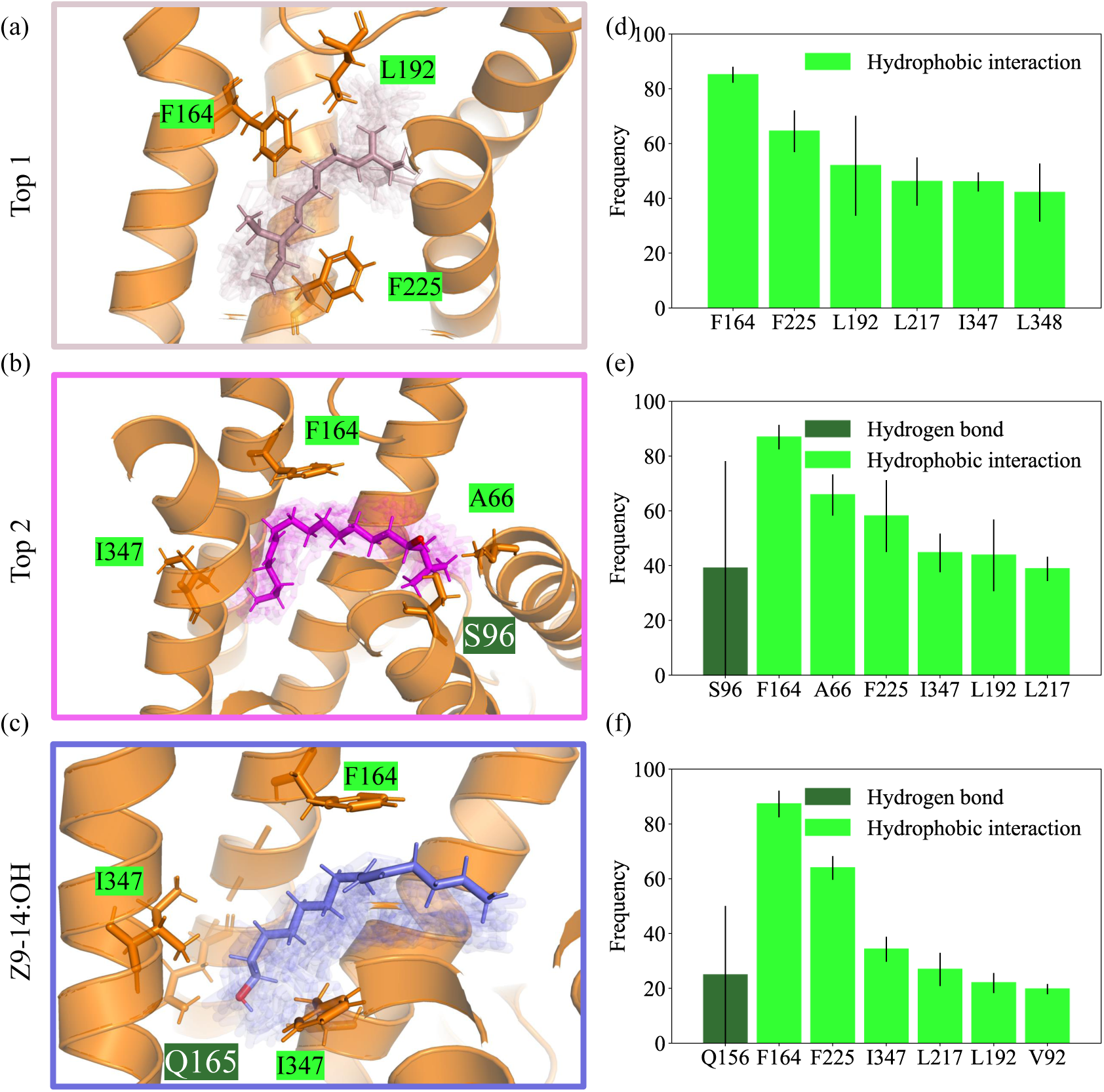
Conformations and contact frequency of ligands in all simulation time. (a-d) Representative conformations of Top1, Top2 (as presented on Fig. 2(d)), Z5-14:OH and Z5-10:OAc within the binding pocket. Transparent sticks represent all simulation frames (5 ns intervals). (e-h) High-frequency interacting residues of SfruOR16 with Top1, Top2, Z9-14:OH respectively. Only the top six hydrophobic and hydrogen bond interactions occurring in more than 20% of frames are shown.

**Table S1:** The CID, IUPAC names, and SIMLES of 49 pheromones obtained from iORbase and previous reports.

| No. | CID | IUPAC name | SMILES |
| --- | --- | --- | --- |
| 1 | 10408 | 6,10,14-trimethylpentadecan-2-one | CC(=O)CCCC(C)CCCC(C)CCCC(C)C |
| 2 | 11005 | tetradecanoic acid | CCCCCCCCCCCCCCCC(=O)O |
| 3 | 11745660 | (E)-3,7-dimethyloct-2-enedioic acid | CC(=CC(=O)O)CCCC(C)C(=O)O |
| 4 | 193373 | 2,6-dimethyloctanedioic acid | CC(CCCC(C)C(=O)O)CC(=O)O |
| 5 | 2266 | nonanedioic acid | O=C(O)CCCCCCCC(=O)O |
| 6 | 5283387 | (Z)-octadec-9-enamide | CCCCCCCCC=CCCCCCCCC(N)=O |
| 7 | 5284421 | methyl (9Z,12Z)-octadeca-9,12-dienoate | CCCCC=CCC=CCCCCCCCC(=O)OC |
| 8 | 5312457 | (2E,4E)-octadeca-2,4-dienoic acid | CCCCCCCCCCCCCCC=CC=CC(=O)O |
| 9 | 5463875 | [(E)-oct-2-enyl] acetate | CCCCC=CCOC(C)=O |
| 10 | 637090 | (E)-but-2-enoic acid | CC=CC(=O)O |
| 11 | 69421 | hexadecanamide | CCCCCCCCCCCCCCCCC(N)=O |
| 12 | 8181 | methyl hexadecanoate | CCCCCCCCCCCCCCCCC(=O)OC |
| 13 | 985 | hexadecanoic acid | CCCCCCCCCCCCCCCCC(=O)O |
| 14 | 984 | hexadecanal | CCCCCCCCCCCCCCCCC=O |
| 15 | 5364495 | (Z)-hexadec-11-enal | CCCCC=CCCCCCCCCCCC=O |
| 16 | 10922550 | (10E,12Z)-hexadeca-10,12-dienal | CCCC=CC=CCCCCCCCCCC=O |
| 17 | 6436633 | [(9E,12Z)-tetradeca-9,12-dienyl] acetate | CC=CCC=CCCCCCCCCOC(C)=O |
| 18 | 5363285 | [(Z)-hexadec-9-enyl] acetate | CCCCCCC=CCCCCCCCCOC(C)=O |
| 19 | 5367650 | [(E)-tetradec-11-enyl] acetate | CCC=CCCCCCCCCCCCOC(C)=O |
| 20 | 5364497 | (Z)-octadec-13-enal | CCCCC=CCCCCCCCCCCCC=O |
| 21 | 6441057 | [(9Z,11E)-tetradeca-9,11-dienyl] acetate | CCC=CC=CCCCCCCCCOC(C)=O |
| 22 | 6436736 | (9E,12Z)-tetradeca-9,12-dien-1-ol | CC=CCC=CCCCCCCCCO |
| 23 | 5364714 | [(Z)-tetradec-9-enyl] acetate | CCCCC=CCCCCCCCCOC(C)=O |
| 24 | 5364471 | (Z)-tetradec-9-enal | CCCCC=CCCCCCCCC=O |
| 25 | 5364643 | (Z)-hexadec-9-enal | CCCCCCC=CCCCCCCCC=O |
| 26 | 5364712 | (Z)-tetradec-9-en-1-ol | CCCCC=CCCCCCCCCO |
| 27 | 5364711 | [(Z)-hexadec-11-enyl] acetate | CCCCC=CCCCCCCCCCCCOC(C)=O |
| 28 | 5367661 | (Z)-hexadec-9-en-1-ol | CCCCCCC=CCCCCCCCCO |
| 29 | 5283305 | (Z)-hexadec-11-en-1-ol | CCCCC=CCCCCCCCCCCCO |
| 30 | 5367599 | [(Z)-tetradec-9-enyl] formate | CCCCC=CCCCCCCCCOC=O |
| 31 | 5363513 | [(Z)-dec-5-enyl] acetate | CCCCC=CCCCCOC(C)=O |
| 32 | 12531 | tetradecyl acetate | CCCCCCCCCCCCCCCCOC(C)=O |
| 33 | 6437763 | (9E,11Z)-tetradeca-9,11-dien-1-ol | CCC=CC=CCCCCCCCCO |
| 34 | 5363405 | [(Z)-dodec-9-enyl] acetate | CCC=CCCCCCCCCOC(C)=O |
| 35 | 5363527 | [(Z)-dodec-7-enyl] acetate | CCCCC=CCCCCCCCOC(C)=O |
| 36 | 5364438 | (Z)-hexadec-7-enal | CCCCCCCCC=CCCCCCC=O |
| 37 | 5367692 | [(Z)-tetradec-11-enyl] acetate | CCC=CCCCCCCCCCCCOC(C)=O |
| 38 | 445639 | (Z)-octadec-9-enoic acid | CCCCCCCCC=CCCCCCCCC(=O)O |
| 39 | 8205 | dodecyl acetate | CCCCCCCCCCCCCOC(C)=O |
| 40 | 69968 | octadecyl acetate | CCCCCCCCCCCCCCCCCCCCOC(C)=O |
| 41 | 12393 | hexadecyl acetate | CCCCCCCCCCCCCCCCCOC(C)=O |
| 42 | 12565 | 6,7-dimethoxy-2,2-dimethylchromene | COC1=C(OC)C=C2OC(C)(C)C=CC2=C1 |
| 43 | 445128 | (10E,12Z)-hexadeca-10,12-dien-1-ol | CCCC=CC=CCCCCCCCCO |
| 44 | 1787910 | (8E,10E)-dodeca-8,10-dien-1-ol | CC=CC=CCCCCCCCO |
| 45 | 5281162 | ethyl (2E,4Z)-deca-2,4-dienoate | CCCCC=CC=CC(=O)OCC |
| 46 | 5363237 | [(E)-dodec-8-enyl] acetate | CCCC=CCCCCCCCCOC(C)=O |
| 47 | 5363377 | [(Z)-dodec-8-enyl] acetate | CCCC=CCCCCCCCCOC(C)=O |
| 48 | 5283295 | (Z)-dodec-8-en-1-ol | CCCC=CCCCCCCCO |
| 49 | 1747818 | [(8E,10E)-dodeca-8,10-dienyl] acetate | CC=CC=CCCCCCCCCOC(C)=O |

**Table S2:**
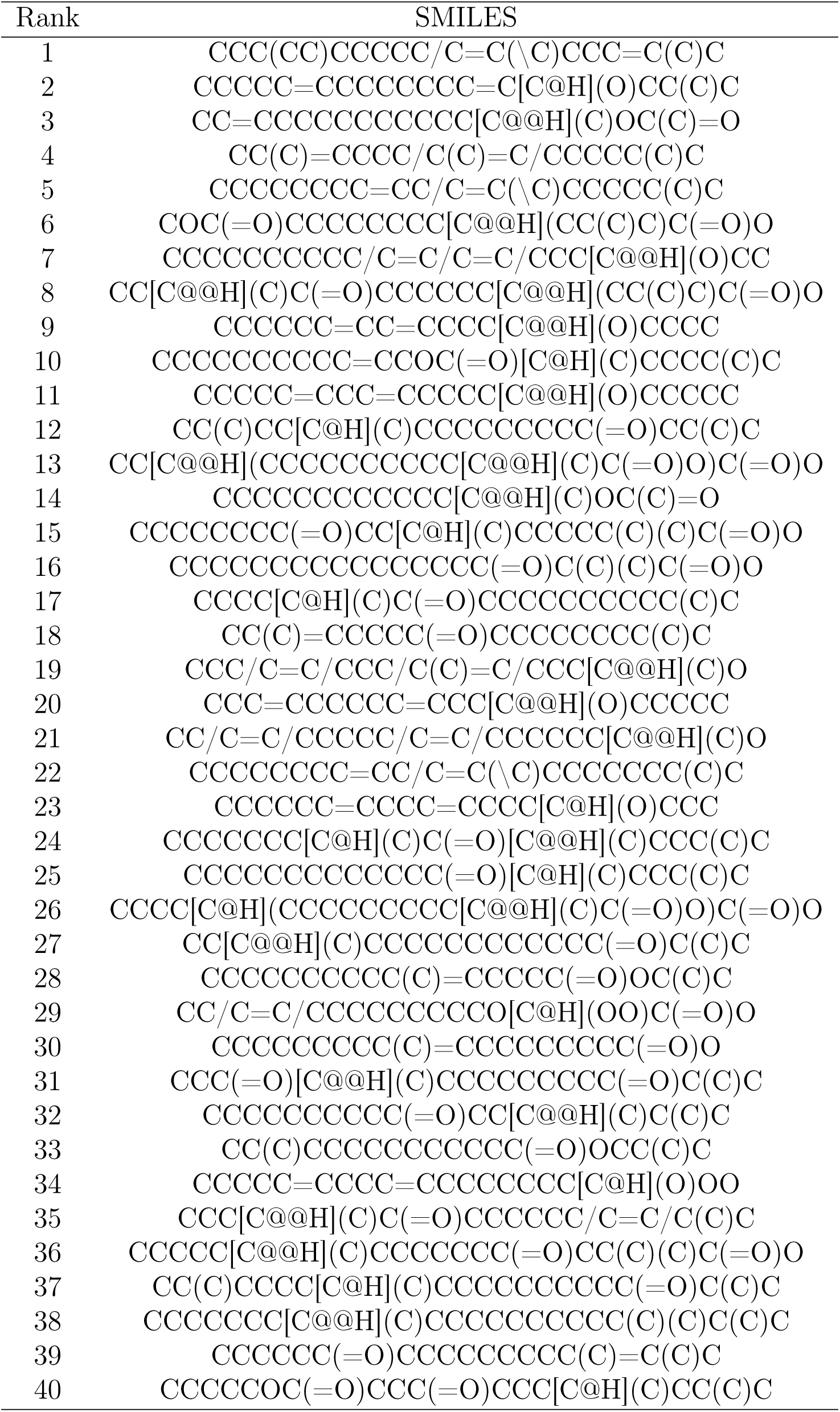
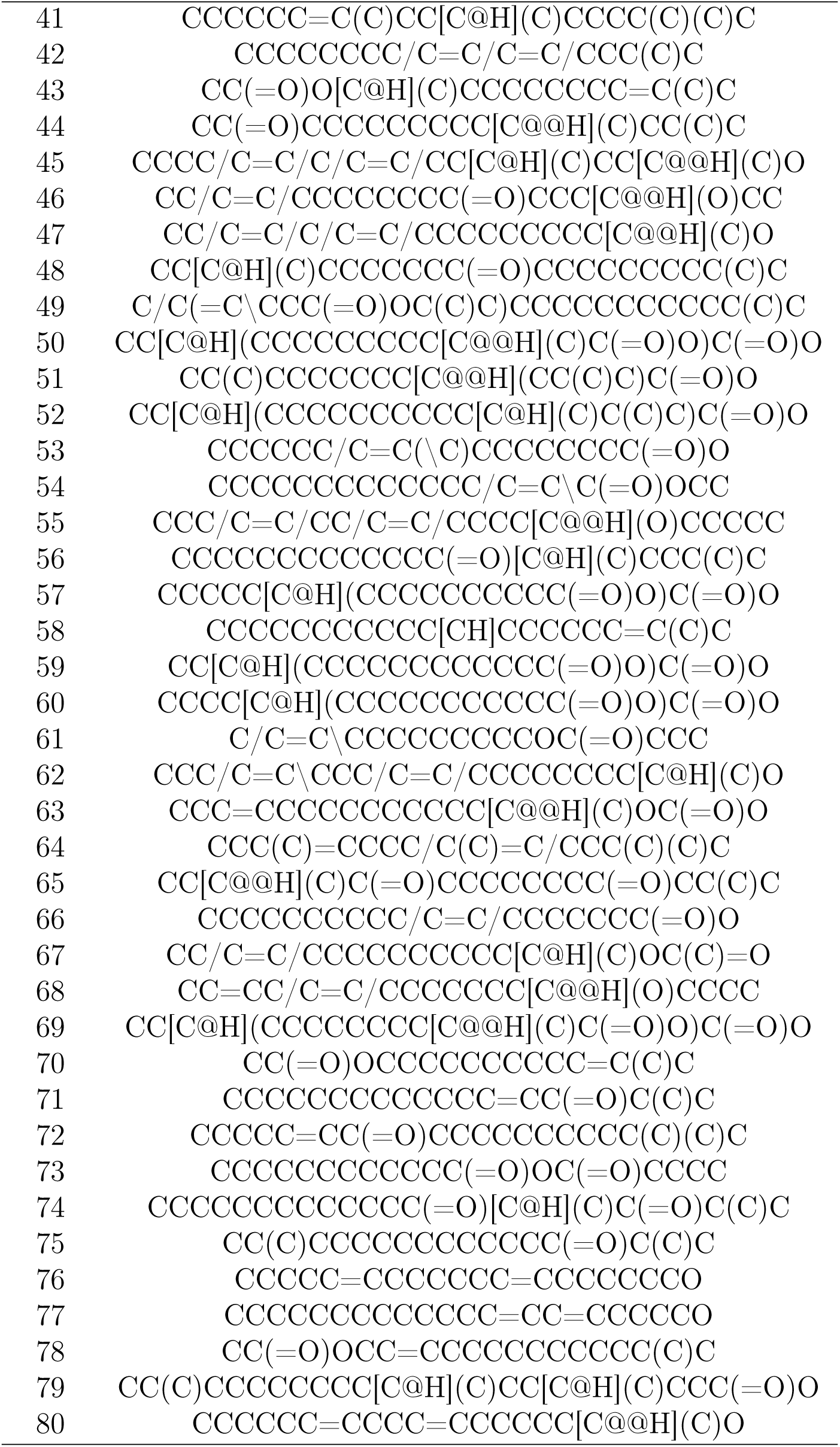
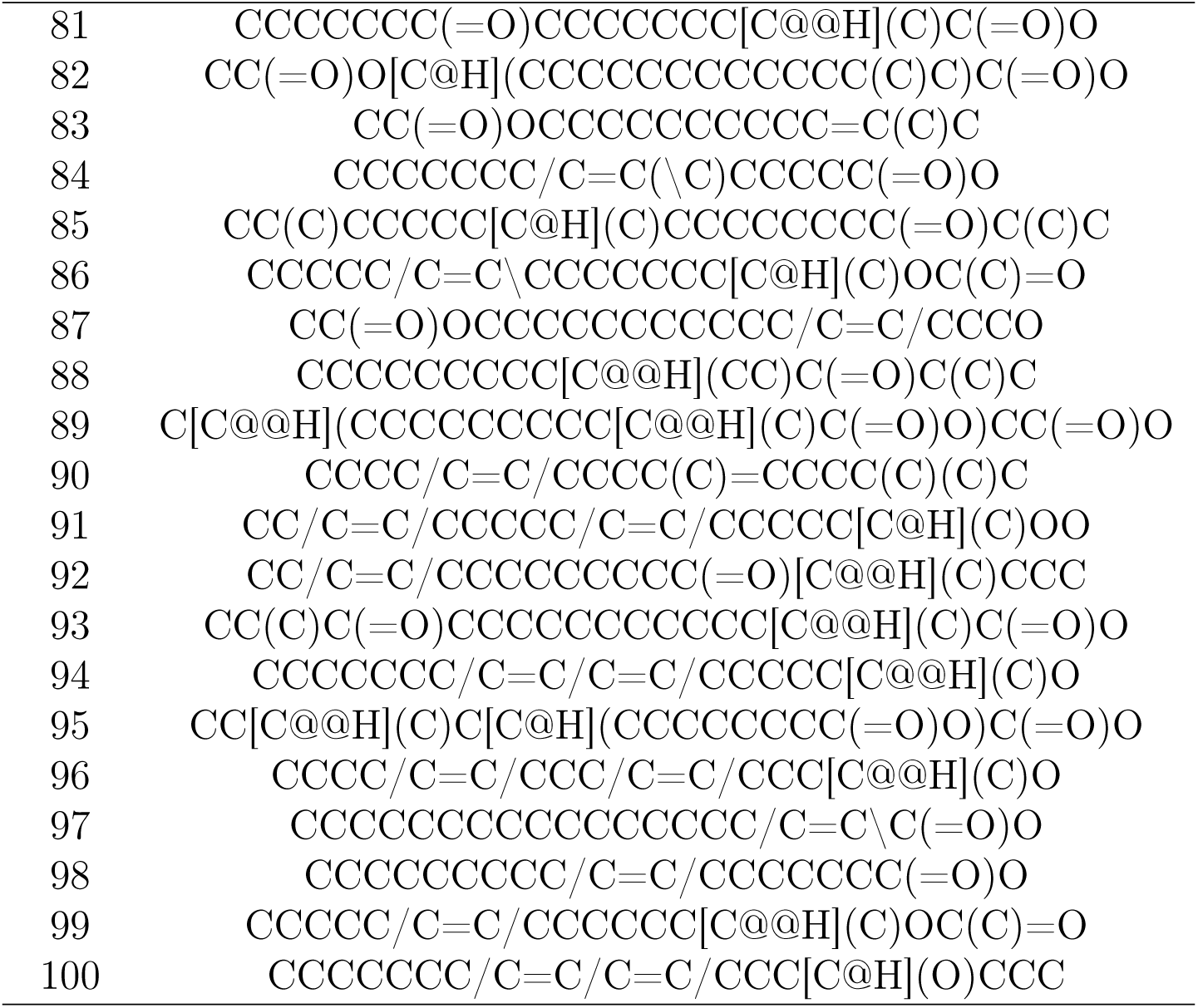
The SIMLES of top 100 sex pheromone analogs generated by GAN after reweighting three rounds corresponding to. Figure S1.

## References

(1) Eberhardt, J.; Santos-Martins, D.; Tillack, A. F.; Forli, S. AutoDock Vina 1.2.0: New docking methods, expanded force field, and Python bindings. J. Chem. Inf. Model. 2021, 61, 3891–3898.

(2) Tang, S.; Chen, R.; Lin, M.; Lin, Q.; Zhu, Y.; Ding, J.; Hu, H.; Ling, M.; Wu, J. Accelerating AutoDock Vina with GPUs. Molecules 2022, 27, 3041.

(3) Cheng, Y.; Gong, Y.; Liu, Y.; Song, B.; Zou, Q. Molecular design in drug discovery: a comprehensive review of deep generative models. Brief. Bioinform. 2021, 22, bbab344.

(4) Zhang, O.; Wang, T.; Weng, G.; Jiang, D.; Wang, N.; Wang, X.; Zhao, H.; Wu, J.; Wang, E.; Chen, G.; Deng, Y.; Pan, P.; Kang, Y.; Hsieh, C.-Y.; Hou, T. Learning on topological surface and geometric structure for 3D molecular generation. Nat. Comput. Sci. 2023, 3, 849–859.

(5) National Library of Medicine (US) PubChem. https://pubchem.ncbi.nlm.nih.gov/, 2004.

(6) Irwin, J. J.; Tang, K. G.; Young, J.; Dandarchuluun, C.; Wong, B. R.; Khurelbaatar, M.; Moroz, Y. S.; Mayfield, J.; Sayle, R. A. ZINC20-A free ultralarge-scale chemical database for ligand discovery. J. Chem. Inf. Model. 2020, 60, 6065–6073.

(7) Böhm, H. J. LUDI: rule-based automatic design of new substituents for enzyme inhibitor leads. J. Comput. Aided Mol. Des. 1992, 6, 593–606.

(8) Xie, Y.; Shi, C.; Zhou, H.; Yang, Y.; Zhang, W.; Yu, Y.; Li, L. {MARS}: Markov Molecular Sampling for Multi-objective Drug Discovery. Proc. Int. Conf. Learn. Represent. 2021.

(9) Bieniek, M. K.; Cree, B.; Pirie, R.; Horton, J. T.; Tatum, N. J.; Cole, D. J. An open-source molecular builder and free energy preparation workflow. Commun. Chem. 2022, 5, 136.

(10) Sánchez-Lengeling, B.; Outeiral, C.; Guimaraes, G.; Aspuru-Guzik, A. Optimizing distributions over molecular space. An Objective-Reinforced Generative Adversarial Network for Inverse-design Chemistry (ORGANIC). ChemRxiv 5309668.v3.

(11) Green, H.; Koes, D. R.; Durrant, J. D. DeepFrag: a deep convolutional neural network for fragment-based lead optimization. Chem. Sci. 2021, 12, 8036–8047.

(12) Imrie, F.; Bradley, A. R.; van der Schaar, M.; Deane, C. M. Deep generative models for 3D linker design. J. Chem. Inf. Model. 2020, 60, 1983–1995.

(13) Zheng, S.; Lei, Z.; Ai, H.; Chen, H.; Deng, D.; Yang, Y. Deep scaffold hopping with multimodal transformer neural networks. J. Cheminform. 2021, 13, 87.

(14) Scott, O. B.; Edith Chan, A. W. ScaffoldGraph: an open-source library for the generation and analysis of molecular scaffold networks and scaffold trees. Bioinformatics 2020, 36, 3930–3931.

(15) Zhang, H.; Zhao, H.; Zhang, X.; Su, Q.; Du, H.; Shen, C.; Wang, Z.; Li, D.; Pan, P.; Chen, G.; Kang, Y.; Hsieh, C.-Y.; Hou, T. Delete: Deep Lead Optimization Enveloped in Protein Pocket through Unified Deleting Strategies and a Structure-aware Network. ArXiv 2023, *abs/2308.02172*.

(16) Goodfellow, I. J.; Pouget-Abadie, J.; Mirza, M.; Xu, B.; Warde-Farley, D.; Ozair, S.; Courville, A.; Bengio, Y. Generative Adversarial Networks. ArXiv 2014, *abs/1406.*2661.

(17) Yu, L.; Zhang, W.; Wang, J.; Yu, Y. SeqGAN: Sequence Generative Adversarial Nets with policy gradient. ArXiv 2016, abs/1609.05473.

(18) Guimaraes, G. L.; Sánchez-Lengeling, B.; Farias, P. L. C.; Aspuru-Guzik, A. Objective-Reinforced Generative Adversarial Networks (ORGAN) for Sequence Generation Models. ArXiv 2017, abs/1705.10843.

(19) Tripathi, S.; Augustin, A. I.; Dunlop, A.; Sukumaran, R.; Dheer, S.; Zavalny, A.; Haslam, O.; Austin, T.; Donchez, J.; Tripathi, P. K.; Kim, E. Recent advances and application of generative adversarial networks in drug discovery, development, and targeting. Artif. Intell. Life Sci. 2022, 2, 100045.

(20) Sato, K.; Pellegrino, M.; Nakagawa, T.; Nakagawa, T.; Vosshall, L. B.; Touhara, K. Insect olfactory receptors are heteromeric ligand-gated ion channels. Nature 2008, 452, 1002–1006.

(21) Cao, S.; Shi, C.; Wang, B.; Xiu, P.; Wang, Y.; Liu, Y.; Wang, G. Evolutionary shifts in pheromone receptors contribute to speciation in four Helicoverpa species. Cell. Mol. Life Sci. 2023, 80, 1–15.

(22) Wang, C.; Cao, S.; Shi, C.; Guo, M.; Sun, D.; Liu, Z.; Xiu, P.; Wang, Y.; Wang, G.; Liu, Y. The novel function of an orphan pheromone receptor reveals the sensory specializations of two potential distinct types of sex pheromones in noctuid moth. Cell. Mol. Life Sci. 2024, 81, 259.

(23) Zhao, J.; Chen, A. Q.; Ryu, J.; Del Mármol, J. Structural basis of odor sensing by insect heteromeric odorant receptors. Science 2024, 384, 1460–1467.

(24) Wang, Y. et al. Structural basis for odorant recognition of the insect odorant receptor OR-Orco heterocomplex. Science 2024, 384, 1453–1460.

(25) Valencia-Montoya, W. A.; Pierce, N. E.; Bellono, N. W. Evolution of sensory receptors. Annu. Rev. Cell Dev. Biol. 2024, 40.

(26) Gu, Z.; Zhao, T.; Li, C.-X. Research progress on odor binding proteins and odor receptors of mosquitoes. Chin. J. Parasitol. Parasit. Dis. 2020, 38, 753.

(27) Zhang, D.-D.; Löfstedt, C. Moth pheromone receptors: gene sequences, function, and evolution. Front. Ecol. Evol. 2015, 3, 00105.

(28) Zhang, S.; Liu, F.; Yang, B.; Liu, Y.; Wang, G.-R. Functional characterization of sex pheromone receptors in Spodoptera frugiperda, S. exigua, and S. litura moths. Insect Sci. 2023, 30, 305–320.

(29) Millar, J. G. Polyene hydrocarbons and epoxides: a second major class of lepidopteran sex attractant pheromones. Annu. Rev. Entomol. 2000, 45, 575–604.

(30) Ando, T.; Inomata, S.-I.; Yamamoto, M. *The Chemistry of Pheromones and Other Semiochemicals I*; Topics in current chemistry; Springer Berlin Heidelberg: Berlin, Heidelberg, 2004; pp 51–96.

(31) Löfstedt, C.; Wahlberg, N.; Millar, J. G. In Pheromone Communication in Moths; Allison, J. D., Carde, R. T., Eds.; University of California Press: Berkeley, 2019; pp 43–78.

(32) Grant, G. G.; Millar, J. G.; Trudel, R. Pheromone identification of Dioryctria abietivorella (Lepidoptera: Pyralidae) from an eastern North American population: geographic variation in pheromone response. Can. Entomol. 2009, 141, 129–135.

(33) Cao, C.; Cao, Y.; Wang, G.-R. Research progress of pheromone receptors in moths. Acta. Entomol. Sin. 2020, 63, 1564–1568.

(34) Li, Q.; Zhang, Y.-F.; Zhang, T.-M.; Wan, J.-H.; Zhang, Y.-D.; Yang, H.; Huang, Y.; Xu, C.; Li, G.; Lu, H.-M. iORbase: A database for the prediction of the structures and functions of insect olfactory receptors. Insect Sci. 2023, 30, 1245–1254.

(35) O’Boyle, N. M. Towards a Universal SMILES representation - A standard method to generate canonical SMILES based on the InChI. J. Cheminform. 2012, 4, 22 –22.

(36) Weininger, D. SMILES, a chemical language and information system. 1. Introduction to methodology and encoding rules. J. Chem. Inf. Comput. Sci. 1988, 28, 31–36.

(37) Landrum, G. RDKit: Open-Source Cheminformatics. https://www.rdkit.org/, 2017.

(38) Yang, L.; Zhang, D.; Karniadakis, G. E. Physics-informed generative adversarial networks for stochastic differential equations. ArXiv 2018, *abs/1811.02033*.

(39) Jumper, J. et al. Highly accurate protein structure prediction with AlphaFold. Nature 2021, 596, 583–589.

(40) Le Guilloux, V.; Schmidtke, P.; Tuffery, P. Fpocket: an open source platform for ligand pocket detection. BMC Bioinformatics 2009, 10, 168.

(41) Jo, S.; Kim, T.; Iyer, V. G.; Im, W. CHARMM-GUI: a web-based graphical user interface for CHARMM. J. Comput. Chem. 2008, 29, 1859–1865.

(42) Lee, J. et al. CHARMM-GUI input generator for NAMD, GROMACS, AMBER, OpenMM, and CHARMM/OpenMM simulations using the CHARMM36 additive force field. J. Chem. Theory Comput. 2016, 12, 405–413.

(43) Huang, J.; Rauscher, S.; Nawrocki, G.; Ran, T.; Feig, M.; de Groot, B. L.; Grubmüller, H.; MacKerell, A. D., Jr CHARMM36m: an improved force field for folded and intrinsically disordered proteins. Nat. Methods 2017, 14, 71–73.

(44) Vanommeslaeghe, K.; Hatcher, E.; Acharya, C.; Kundu, S.; Zhong, S.; Shim, J.; Darian, E.; Guvench, O.; Lopes, P.; Vorobyov, I.; Mackerell, A. D., Jr CHARMM general force field: A force field for drug-like molecules compatible with the CHARMM all-atom additive biological force fields. J. Comput. Chem. 2010, 31, 671–690.

(45) Abraham, M. J.; Murtola, T.; Schulz, R.; Páll, S.; Smith, J. C.; Hess, B.; Lindahl, E. GROMACS: High performance molecular simulations through multi-level parallelism from laptops to supercomputers. SoftwareX 2015, 1–2, 19–25.

(46) Valdés-Tresanco, M. S.; Valdés-Tresanco, M. E.; Valiente, P. A.; Moreno, E. Gmx_MMPBSA: A new tool to perform end-state free energy calculations with GROMACS. J. Chem. Theory Comput. 2021, 17, 6281–6291.

(47) Miller, B. R., 3rd; McGee, T. D., Jr; Swails, J. M.; Homeyer, N.; Gohlke, H.; Roitberg, A. E. MMPBSA.Py: An efficient program for end-state free energy calculations. J. Chem. Theory Comput. 2012, 8, 3314–3321.

(48) Guo, J.-M.; Liu, X.-L.; Liu, S.-R.; Wei, Z.-Q.; Han, W.-K.; Guo, Y.; Dong, S.-L. Functional Characterization of Sex Pheromone Receptors in the Fall Armyworm (Spodoptera frugiperda). Insects 2020, 11.

